# Capturing long-tailed individual tree diversity using an airborne multi-temporal hierarchical model

**DOI:** 10.1101/2022.12.07.519493

**Authors:** Ben G. Weinstein, Sergio Marconi, Sarah J Graves, Alina Zare, Aditya Singh, Stephanie A Bohlman, Lukas Magee, Daniel J. Johnson, Phillip A. Townsend, Ethan P. White

## Abstract

Measuring forest biodiversity using terrestrial surveys is expensive and can only capture common species abundance in large heterogeneous landscapes. In contrast, combining airborne imagery with computer vision can generate individual tree data at the scales of hundreds of thousands of trees. To train computer vision models, ground-based species labels are combined with airborne reflectance data. Due to the difficulty of finding rare species in a large landscape, the majority of classification models only include the most abundant species, leading to biased predictions at broad scales. Extending classification models to include rare species requires targeted data collection and algorithmic improvements to overcome large data imbalances between dominant and rare taxa. In addition, large landscapes often require multiple acquisition events, leading to significant within-species variation in reflectance spectra. Using a multi-temporal hierarchical model, we demonstrate the ability to include species predicted at less than 1% frequency in landscape without losing performance on the dominant species. The final model has over 75% accuracy for 14 species with improved rare species classification compared to a baseline deep learning model. After filtering out dead trees, we generate landscape species maps of individual crowns for over 670,000 individual trees at the Ordway Swisher Biological Station within the National Ecological Observatory Network. We estimate the relative abundance of the species within the landscape and provide three measures of uncertainty to generate a range of counts for each species. These maps provide the first estimates of canopy tree diversity within NEON sites to include rare species and provide a blueprint for capturing tree diversity using airborne computer vision at broad scales.

## Introduction

Forest ecosystem services, ecology, and biogeography all depend on the composition of individual tree species. However, traditional methods for collecting species information cannot scale to entire forests and are mostly limited to sampling plots with small spatial coverage. In contrast, airborne data collection using hyperspectral sensors can cover areas containing millions of trees. Early studies of remotely-sensed tree species prediction focused largely on hand-crafted features using spectral indices to create machine learning models (e.g. Heikkinen et al. 2010, Schäfer et al. 2016, Maschler et al. 2018). These studies often used coarse taxonomic categories (e.g. ‘conifer’) and were restricted to a small number of bands across a wide spectral range (Dalponte and Coomes 2016). Following the trend away from hand-crafted features towards deep learning neural networks, numerous publications have applied computer vision approaches to airborne tree species prediction (Mäyrä et al. 2021, Abbas et al. 2021, Chen et al. 2022, Onishi et al. 2022, Veras et al. 2022). These applications demonstrated that models often achieve between 70% to 90% accuracy for fewer than 10 co-occurring, well-sampled, classes (Marconi et al. 2022). A remaining species classification challenge is broadening classifications beyond the most common and differentiable species and applying these models to the scale of entire forested landscapes. Most previous work focused on either relatively species poor areas, (e.g. four species in Mäyrä et al. 2021), or only included the most common species in small sampling areas (Veras et al. 2022). Extending the number of classes in a species classification model broadens the questions that can be answered using airborne remote sensing to include biodiversity modeling and fine-grained, wildlife habitat management. Additionally, we can be more confident in predicted tree crown map accuracy since they are more likely to cover the species in the landscape. However, as species number increases, species misclassifications also increase, making it more difficult to learn discriminative features (Qin et al. 2022). This challenge is further compounded when applying species classification models to large areas. For example, Onishi et al. (2022) showed that the accuracy of an RGB deep learning tree species model dropped by more than 35% among acquisition flights.

Our goal was to develop an individual tree species model that captures common and rare tree species across a 3800 hectare landscape at the Ordway-Swisher BIological Station, a National Ecological Observatory Network (NEON) site in Florida. NEON includes 47 terrestrial sites across the United States, with vegetation data in form of forest plots in which individual trees are identified to species and overlapping airborne LIDAR, RGB and hyperspectral data sampled annually for most sites. Airborne species classification within NEON sites started with Fricker et al. (2019) at the Teakettle California site (TEAK) using 7 species and a ‘Dead’ class collected during one flightline. Scholl et al. (2020) followed with a four species model at the Niwot Ridge NEON site in Colorado (NIWO). Two data science competitions, one focused on the OSBS site (Marconi et al. 2019) and the second combining data from the OSBS, Talladega, Alabama (TALL), and Mountain Lake, Virginia (MLBS) sites in the Southeastern US (Graves et al. 2021), used NEON forest plot data with 33 species, of which only 15 had more than 5 individuals. The lack of data for less common species was the primary factor in poor model performance. For example, as part of the multi-site data competition (Graves et al. 2021), Scholl et al. (2021) modeled 27 selected species classes, but only 7 of these classes had non-zero evaluation accuracy. Marconi et al. (2021) attempted the first NEON-wide model for 77 species across 27 sites using a pixel-based ensemble machine learning classifier. In all of these studies, only the most common species were classified, largely due to insufficient field data on rare species for model development and evaluation. This illustrates that even well-designed vegetation sampling (NEON’s forest plots are stratified by habitat type to capture landscape variation) that is used for developing remote sensing models often lacks sufficient field samples of rare species, requiring approaches to supplementing available data and building models that are robust to small amounts of data in rare classes.

While the number of rare species naturally differs across ecosystems, our goal was to develop modeling approaches that can be applied to the rarest species and be used to capture diversity of ecosystems with canopy trees with long tails of rare species. In the case of the OSBS NEON forest plots contain only five canopy species with more than five individuals; however, we estimate that there are at least 18 canopy tree species across the landscape. This difference is in part due to NEON’s plot design which focuses on covering the common land types in the ecosystem using the National Land Cover Database. While we know five species dramatically underestimate tree diversity at the site, it is not yet possible to estimate the proportion of crowns they represent. Using fixed-plot data biases samples towards common species and underrepresents the role of rare species at landscape levels, while using targeted sampling overrepresents rare species. Airborne remote sensing provides a more unbiased estimate of the tree community composition at the forest scale. However, there are limitations to the maximum species diversity that can be captured in airborne classification models. Species with only a handful of individuals are unlikely to generalize across broad areas, even with targeted data collection.

Broadening the number of species in remote sensing classification models to include rarer species creates several challenges. The largest problem is sample imbalance between common and rare species. Machine learning models favor predictions that improve overall performance and often ignore rare species classes. Due to small sample sizes, we risk overfitting models applied to spectra from individuals of rare species. To combat both the risk of class imbalance and rare species overfitting, we developed a hierarchical, multi-temporal ensembling approach using convolutional neural networks with spectral attention. The spectral attention blocks act as a form of data reduction to help the model learn which combinations of bands are most useful in classification. This reduces the overfitting by reducing hundreds of spectral bands into a narrower number of classification features. The multi-temporal aspect uses repeated views of the same ground truth tree across years to bolster low sample sizes of rare species. The learned features are also more robust to annual variation in spectral reflectance that arises through differences in sampling conditions and the data acquisition environment. Finally, the hierarchical model combines multiple sub-models that each distinguish only a few classes. This combination helps address the inherent class imbalance in species abundances and the challenge of developing discriminative features with many classes, while still avoiding undersampling common classes or oversampling rare classes.

Using a hierarchical multi-temporal model, we generated landscape level maps for hundreds of thousands of individual tree crowns. Using these maps, we estimated species abundance distributions at broad scales. However, this biodiversity point estimate did not incorporate any uncertainty in the predicted species counts. It did not indicate the relative confidence among classes, and it provided no way of assessing prediction accuracy. To address these shortcomings, we assessed the different types of uncertainty from our species predictions. There was uncertainty in data due to trees selected as training versus evaluation examples during model development, in the model due to the stochastic nature of learning features for classification, and in each predicted crown through the model's confidence score. By evaluating these areas of uncertainty we begin to bring deep learning in ecological monitoring closer to traditional monitoring programs that have robust confidence measures. While these approaches to capture rare species and describing uncertainty were tested at one site, we expect these approaches to be broadly useful at any site with long-tailed tree diversity.

## Methods

We developed a multi-stage pipeline to predict tree species based on field-collected labels and airborne sensor data (Figure 1). In summary, researchers collected crown data and identified tree species in the field using a geospatial point at the tree stem, or drew a tree crown on a tablet while viewing the corresponding airborne imagery. The stem data were converted into a crown using an existing RGB tree crown deep learning model (Weinstein et al. 2020a). Tree crown locations were then judged to be visible from the air by comparing LiDAR-derived heights and field measured heights, which were available only for NEON field plot data. Hyperspectral data were cropped to each tree crown, split into training and test segments, and used to train a multi-temporal hierarchical model. During prediction, the same RGB tree crown deep learning model was used to generate predicted crowns, which were then assigned a species label using hyperspectral data. This workflow was applied to four years of airborne data and all tiles within the bounds of the OSBS NEON site to create maps of hundreds of thousands of individual trees.

**Figure 1.**
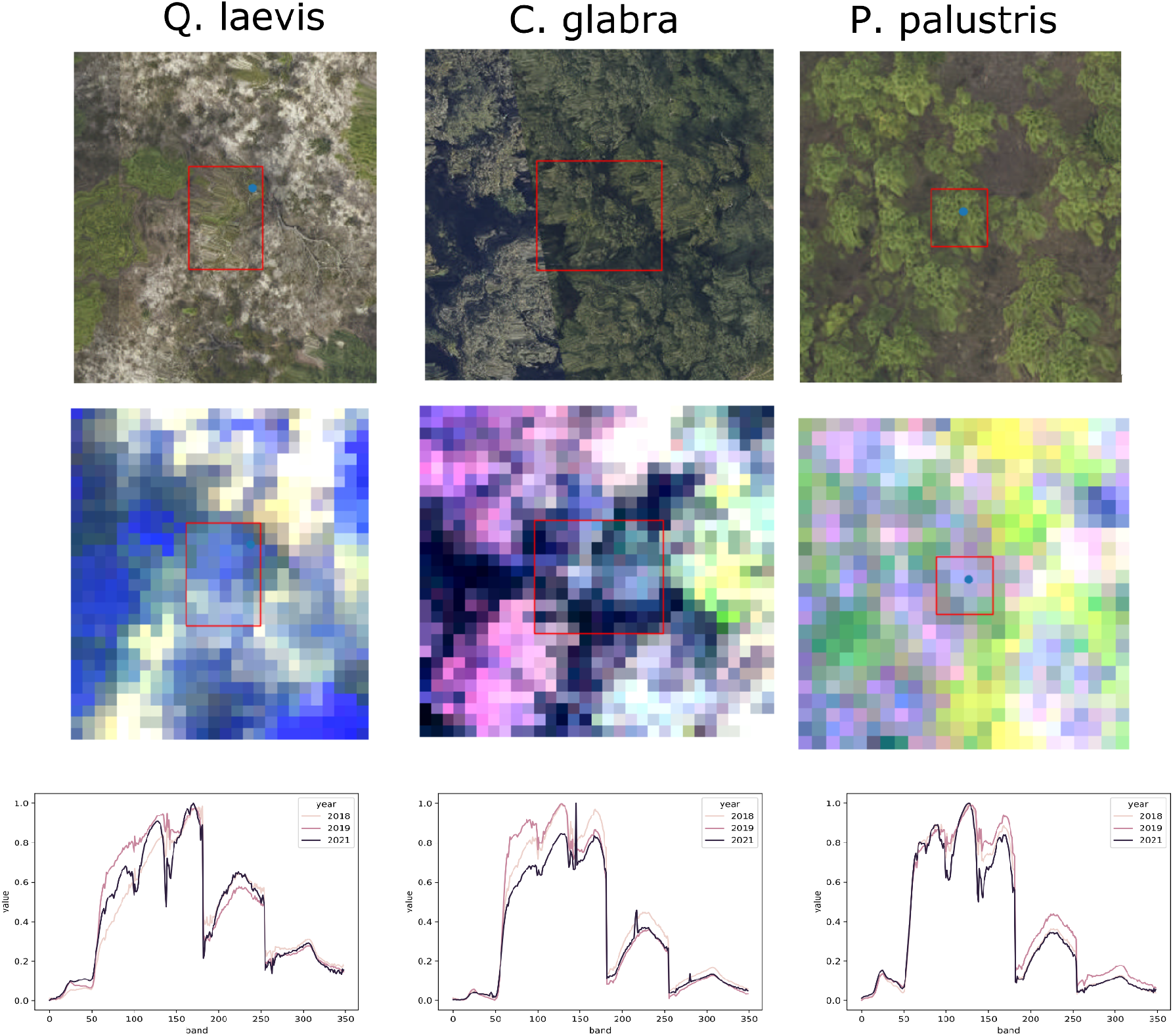
Input data and sensor data types for three sample species. Crown locations (red bounding box) are either estimated directly in the field using a tablet (top-center), or predicted from a GPS point at the tree location (blue point) using a deep learning RGB model. These data are overlaid on 369 hyperspectral data bands, shown here as a three-band false color image (457 nm, 648 nm, 939 nm wavelengths). The spectral signature for each sample is collected for each available year of sensor data.

### Data Collection

NEON collected airborne data over the Ordway-Swisher Biological Station annually in September between 2017 and 2021, with the exclusion of 2020 due to pandemic-related closures. We used four data products from NEON’s airborne observation platform, 1) orthorectified Camera Mosaic (‘RGB’ NEON ID: DP3.30010.001), 2) ecosystem sStructure (‘Canopy Height Model’ NEON ID: DP3.30015.001), 3) hyperspectral surface reflectance (‘HSI’ NEON ID: DP1.30006.001), and 4) vegetation structure (NEON ID: DP1.10098.001) (National Ecological Observatory Network (NEON) 2021). The 10 cm RGB data were used to predict tree crown locations necessary for associating field labels and sensor data during model development. RGB data were also used to identify dead trees during our prediction workflow. The 1 m canopy-height model was used to determine which field collected data were likely to be visible from the air, as well as to define a 3-m minimum tree height threshold during the prediction workflow. The HSI data is used to differentiate tree species based on spectral reflectance. The HSI data spanned approximately 420-2500 nm with a spectral sampling interval of 5 nm, a total of 426 bands. NEON provides orthorectified images with a pixel size of 1 m^2^ in 1 km^2^ tiles that are georectified and aligned with the RGB and Canopy-Height-Model. For more information on hyperspectral data processing and calibration see NEON technical document NEON.DOC.001288.

The NEON Vegetation Structure dataset is a collection of tree stem points within fixed-width field plots; plot locations are allocated across sites according to a stratified random, spatially balanced design (Barnett et al. 2019). The plot locations were designed to capture the major ecosystems within a site. All trees in sampled areas with a stem diameter > 10 cm are mapped and measured for diameter and height, and health status and species identity are recorded. Building on this NEON dataset, we incorporated stem location and information from Wang et al. (2020) and the OSBS ForestGEO plot (Johnson et al. 2021). We targeted sampling rare or incorrectly predicted species using preliminary species classification results and identifying ecosystem types with high frequency of rarer species from the Florida Natural Areas Inventory (fnai.org) of the site (Figure S1). We used tree crown locations predicted from Weinstein et al. (2021) to identify target trees in the field and map them on the images following Graves et al. (2020), resulting in 1556 additional trees from 24 areas of interest. Several rare species were found but discarded for the following reasons:1) any species with less than five individuals found during field sampling (n=5), 2) any shrub-like species judged not to be visible in the canopy (e.g. *Ilex cassine)*, 3) species whose taxonomic status was in question, e.g. *Quercus laurifolia* versus *Quercus hemisphaerica*. The final number of samples per species is shown in Table 1.

**Table 1.**
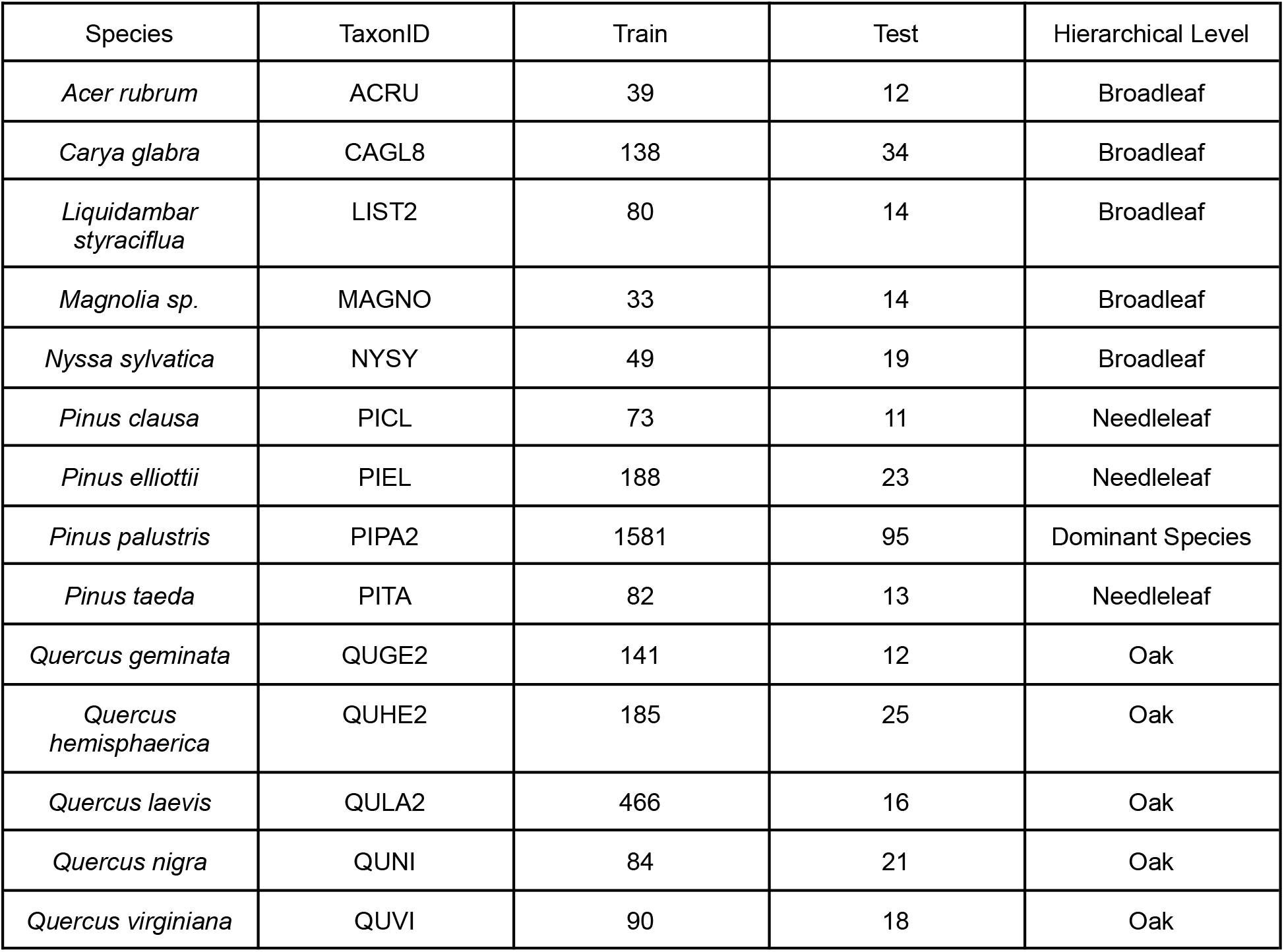
The number of training and test individuals for each taxa. The NEON taxonomic identification codes are used throughout the figures for abbreviations. Hierarchical level indicates how the species was classified in the hierarchical model.

| Species | TaxonID | Train | Test | Hierarchical Level |
| --- | --- | --- | --- | --- |
| *Acer rubrum* | ACRU | 39 | 12 | Broadleaf |
| *Carya glabra* | CAGL8 | 138 | 34 | Broadleaf |
| *Liquidambar styraciflua* | LIST2 | 80 | 14 | Broadleaf |
| *Magnolia sp.* | MAGNO | 33 | 14 | Broadleaf |
| *Nyssa sylvatica* | NYSY | 49 | 19 | Broadleaf |
| *Pinus clausa* | PICL | 73 | 11 | Needleleaf |
| *Pinus elliotii* | PIEL | 188 | 23 | Needleleaf |
| *Pinus palustris* | PIPA2 | 1581 | 95 | Dominant Species |
| *Pinus taeda* | PITA | 82 | 13 | Needleleaf |
| *Quercus geminata* | QUGE2 | 141 | 12 | Oak |
| *Quercus hemisphaerica* | QUHE2 | 185 | 25 | Oak |
| *Quercus laevis* | QULA2 | 466 | 16 | Oak |
| *Quercus nigra* | QUNI | 84 | 21 | Oak |
| *Quercus virginiana* | QUVI | 90 | 18 | Oak |

To connect airborne sensor data with species labels collected in the field, we converted individual stems into a tree crown using either the bounding box of the field-drawn polygon or the field stem point. For the field stem points, we used the DeepForest model to predict bounding boxes that overlapped the field points. The DeepForest model is a RGB tree detection tool that has been validated using NEON data (Weinstein et al. 2020a, 2020b, 2021). While previous work used a fixed number of pixels closest to the field stem location (Scholl et al. 2020, Marconi et al. 2022), model-based bounding boxes linked training and evaluation processes more directly to predictions where no stem locations were available. In the event that multiple predicted tree crown boxes overlapped the field point, we chose the box centroid nearest the field stem.

Due to differences in data acquisition timing and sampling conditions, raw reflectance values differed among flight years. We found that the optimal normalization strategy divided the value of each hyperspectral channel by the maximum value for that channel across the entire crown. To create a consistent input size, we resized the crop to 11 × 11 pixels using nearest neighbors interpolation. Several reasonable sizes were tested without considerable variation. We removed bands associated with water absorption, which were usually completely saturated and non-informative for tree species prediction. We also removed the first and last 10 bands due sensor noise, resulting in a total of 349 bands as model inputs.

## Modeling Approaches

To build a multi-temporal hierarchical model, we tested a series of submodels to assess overall model performance at each progressive stage. We began with a 2D convolutional neural network model, which is the most commonly used model architecture in computer vision. We then added a depth-wise spectral attention architecture to customize the 2D CNN to hyperspectral data. These 2D spectral attention models were then stacked to form a multi-temporal ensemble. We next organized the multi-temporal ensembles into a hierarchical structure that separately modeled individual subgroups.

### 2D CNN

The baseline model was a set of two-dimensional convolutional neural blocks (2D CNN). Each 2D CNN block has a kernel size of 3 × 3, followed by 2D batch normalization and RELU activation. The 2D CNN model has three 2D CNN blocks with progressively deeper filters of 32, 64, 128 size channels. Max pooling with kernel size 2 × 2 is applied to blocks 2 and 3. To obtain class scores, output from the third block is flattened and passed through a fully connected layer of depth 512 with softmax activation.

### 2D CNN with spectral attention

Building from the 2D CNN model, we added a spectral attention layer to help the model focus on salient combinations of features among HSI bands (Hang et al. 2020). Each spectral attention layer is inserted after each CNN block. The output of each CNN block is pooled across the spatial dimension, creating a 1D feature output. This 1D layer is passed to a 1D convolutional layer. Similar to the 2D CNN block described above, the spectral attention filter is progressively deeper, with 32, 64 and 128 filters and kernel sizes of 3, 5, and 7. Compared to Hang et al. (2020), we removed the spatial attention network, as it provided no benefit and increased overfitting.

### Multi-temporal ensemble

To combat the challenge of within-class spectral variance, we separately modeled 2017, 2018, 2019 and 2021 sensor data followed by an ensemble among years. Using multiple views has the potential to overcome illumination changes among flights and biological variation in leaf signatures due to phenology and health status, thus allowing more robust feature learning for species classification. The ensemble also reduces the potential effect of spatial mismatch between field-collected levels and sensor data, which arise from single-year georectification errors. The final cross-year score is the mean of class scores among years.

### Hierarchical mixture of multi-temporal experts

We organized the multi-temporal ensembles into a nested series of five hierarchical models. The organization is guided by three aims: 1) reduce class imbalance by separately modeling the most common training class and rarer classes, 2) highly similar classes should be grouped together and away from distant classes, allowing submodels to learn features specific to this task, 3) higher level submodels should align with biological groupings, such that downstream users could choose to adopt biological grouped labels over the more refined species specific-labels. The top model classifies a sample as belonging to the most common species, *P. palustris* (PIPA2), or ‘Other.’ If classified as ‘Other,’ a sample is classified as either ‘Broadleaf’ or ‘Needleleaf’. If samples are classified as ‘Needleaf’, they are passed to the Needleleaf submodel for final species classification. If samples are classified as ‘Broadleaf,’ they are passed to a broadleaf submodel that includes all species but lumps ‘Oaks’ into a single class. If predicted ‘Oak,’ the sample is classified by the final oak-only submodel.

### Training

All models were trained on a single NVIDIA A100 GPU with a batch size of 128. Each model was trained for 70 epochs using ADAM optimization and a default learning rate of 0.0001. This learning rate was altered for the hierarchical model for which each submodel had a separate learning rate. We found that higher-order submodels, such as the Broadleaf versus Needleaf level, required a lower learning rate of 0.00001, whereas the lower levels used the default 0.0001 learning rate. The learning rate was reduced every eight epochs by a factor of 0.75 based on decreasing validation loss, with a minimum learning rate of 1e-6. While our hierarchical strategy dramatically reduced class imbalance by grouping rare species, we did provide minimal undersampling of oak classes by allowing no more than 200 samples per species. This is a much higher ceiling than would have been possible without the hierarchical approach. During training, we provided standard rotation and flip augmentation to reduce overfitting. To promote rare species classification, we divided the loss of each sample by species frequency, which has the effect of increased loss weight for rare classes.

### Evaluation

To create a train-test split for model evaluation, we adopted a geographic approach to ensure that data collected in close proximity did not occur in both train and test. For the NEON woody vegetation structure data, entire NEON plots were assigned to either train or test. For the non-NEON data, which did not naturally fall into fixed plots, we drew a 40 × 40 m grid over the entire site and assigned trees within each cell into either train or test. Using this train-test split, we evaluated our species classification models using recall, defined as the proportion of correctly predicted samples, and precision, defined as the proportion of predictions which were correct. We further classified these metrics into micro-averaged and macro-averaged scores. Micro-averaging uses the natural abundance of each class in the test set, whereas macro-averaging first computes the metric on each species individually and then averages the results. Given the large imbalance in the class frequency, the macro-averaged results are sensitive to the rare species performance, whereas the micro-averaged results are sensitive to the dominant species performance.

### Alive-Dead Filtering

One added complexity of this workflow is that our field data set did not contain many dead trees, which regularly occur on the landscape and should be classified separately from living trees. To provide a simple filter for trees that appear dead in the RGB data we collected 6,342 crops from the prediction landscape, as well as other NEON sites, and hand annotated them as either alive or dead. We finetuned a resnet-50 pre-trained on ImageNet to classify alive or dead trees before passing them to the species classification model. The model was trained with an ADAM optimizer with a learning rate of 0.001 and batch size of 128 for 40 epochs, and was evaluated on a randomly held out of 10% of the crops.

### Uncertainty

Our workflow generates counts of each species at the full-site scale. However, this count comes without uncertainty and no mechanism to assess relative confidence among classes or samples. We identified three types of uncertainty in predicted counts: 1) data uncertainty, 2) model uncertainty, and 3) sample-based uncertainty (Hu et al. 2021, Lai et al. 2021, Abdar et al. 2021). Data uncertainty is variance due to individual differences in spectral reflectance. This uncertainty can be measured using k-fold cross validation, where the original pool of data is repeatedly split into random train/test splits, and the entire workflow is repeated, leading to a new set of predicted species counts at the full-site scale. Model uncertainty is variance due to stochastic optimization of model weights. This can be especially important among rare taxa, where the highest predicted class for an individual crown may have an only marginally higher score than the 2nd predicted class. By repeatedly training the network from initialization, we propagated the uncertainty into downstream species predictions. Sample uncertainty is the confidence in each prediction, which is a combination of the model's predicted confidence score, such that we are not assuming all samples are equally likely to be incorrect, as well as the empirical evaluation score that captures which species are likely to be confused. To measure this effect on predictions, we took the output of each sample and assigned the label based on the evaluation confusion matrix. For example, if a sample was predicted to be *Q. geminata* with a confidence score of 70%, we took a Bernoulli random draw with this probability. If the result was zero, we drew a new species prediction from the evaluation confusion for this sample. If *Q. geminata* was correctly classified in 60% of the evaluation samples and confused with *Q. nigra* in 40% of the evaluation samples, the new species ID had a 40% chance of being assigned to *Q. nigra*.

## Results

The hierarchical multi-temporal spectral attention model had the highest performance with 75% micro-accuracy, 64% macro-accuracy over all species, and with 96% precision and recall for the dominant species, *P. palustris* (Table 2). Ten-fold cross-validation of the train-test split showed modest variation in evaluation metrics among train/test splits, but always remained higher than simpler models. Adding spectral attention increased the micro-averaged accuracy of all classes, as well as the precision of the dominant class *P. palustris*, compared to a baseline 2D CNN. The multi-temporal ensemble alone did improve recall of the dominant class to 96%, but at the cost of reduced precision of 87%, meaning that many non-*P. palustris* trees were classified as *P. palustris*. However, the combination of a hierarchical model structure and a multi-temporal ensemble resulted in significant improvements in performance in three out of four metrics (Table 2), indicating an important classification interaction between these two approaches.

**Table 2.** Evaluation scores for each component of the final model. The micro-averaged accuracy is the proportion of samples correctly predicted. The macro-averaged accuracy is the proportion of samples correctly predicted by species, averaged across all species. The recall and precision of the dominant class class, *P. palustris*, are also used as metrics to illustrate the accuracy of species-level predictions and because the dominant species will have a significant impact on the quality of full-site predictions. For the multi-temporal hierarchical model, the training workflow was repeated 10 times beginning from a random initialization of weights to generate a range of evaluation metrics. The mean (min, max) of each metric is shown.

| Model | Micro-accuracy | Macro-accuracy | *P. palustris*<br>Recall | *P. palustris*<br>Precision |
| --- | --- | --- | --- | --- |
| 2D CNN | 0.61 | 0.48 | 0.92 | 0.80 |
| 2D CNN<br>+ Spectral Attention | 0.65 | 0.54 | 0.92 | 0.92 |
| Multi-temporal<br>+ Spectral Attention | 0.66 | 0.52 | 0.96 | 0.87 |
| Hierarchical<br>+ Spectral Attention | 0.60 | 0.59 | 0.91 | 0.91 |
| Multi-temporal +<br>Hierarchical +<br>Spectral Attention | 0.75 (0.71,<br>0.78) | 0.63 (0.58,<br>0.67) | 0.96 (0.95, 0.99) | 0.96 (0.94, 0.98) |

The first model of *P. palustris* versus all other taxa had an accuracy of 97.2% with precision and recall for *P. palustris* of 96.3% and 94.0%, respectively. The second model of ‘Broadleaf’ versus ‘Needleleaf’ had a mean accuracy of 96.4% with higher precision for Broadleaf (99.3%) than Needeleaf (93.6%). The final higher level label is within the ‘Broadleaf’ model which jointly grouped all oak trees. The ‘Oak’ label within this model had a precision of 90.4% and an accuracy of 77.6%. The confusion matrix showed the majority of misclassification occurred in nested models. For example, the most common misclassification of *Quercus geminata* is *Quercus nigra* which both occur within the ‘Oak’ nested model. While there are examples of misclassification across nested models, such as among *Quercus laevis* and *P. palustris* which co-occur in the same habitat, these are relatively infrequent due to the high accuracy of the higher level submodels (Figure 3).

**Figure 2.**
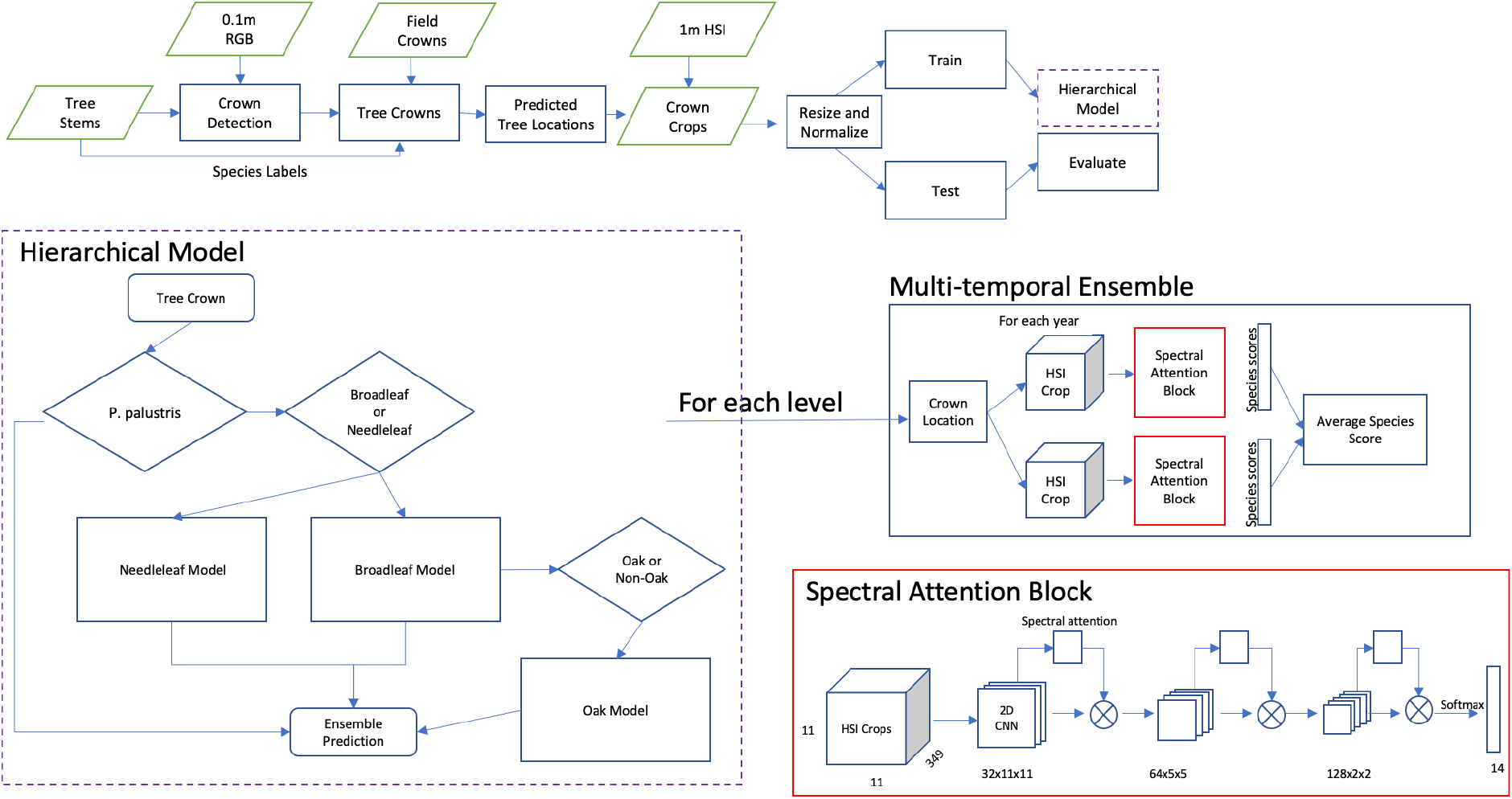
Workflow for classification using a hierarchical multi-temporal model. The field collected data are tree stems or crown polygons. For tree stems, crown locations are detected using a RGB deep learning model (Weinstein et al. 2021). Crown locations were used to crop available sensor data for each year. The cropped tree crowns are resized, normalized, and used to train the hierarchical model. The hierarchical model begins by classifying whether the input sample is the most common class–*P. palustris*. If not classified as *P. palustris*, the broadleaf or needleleaf distinction decides which downstream model is used. The broadleaf model contains a nested category to separate the oak subgroup. For each submodel, we use the same multi-temporal ensemble architecture. Within the multi-temporal ensemble, each year is separately modeled using a spectral attention block. The results are then averaged among years.

**Figure 3.**
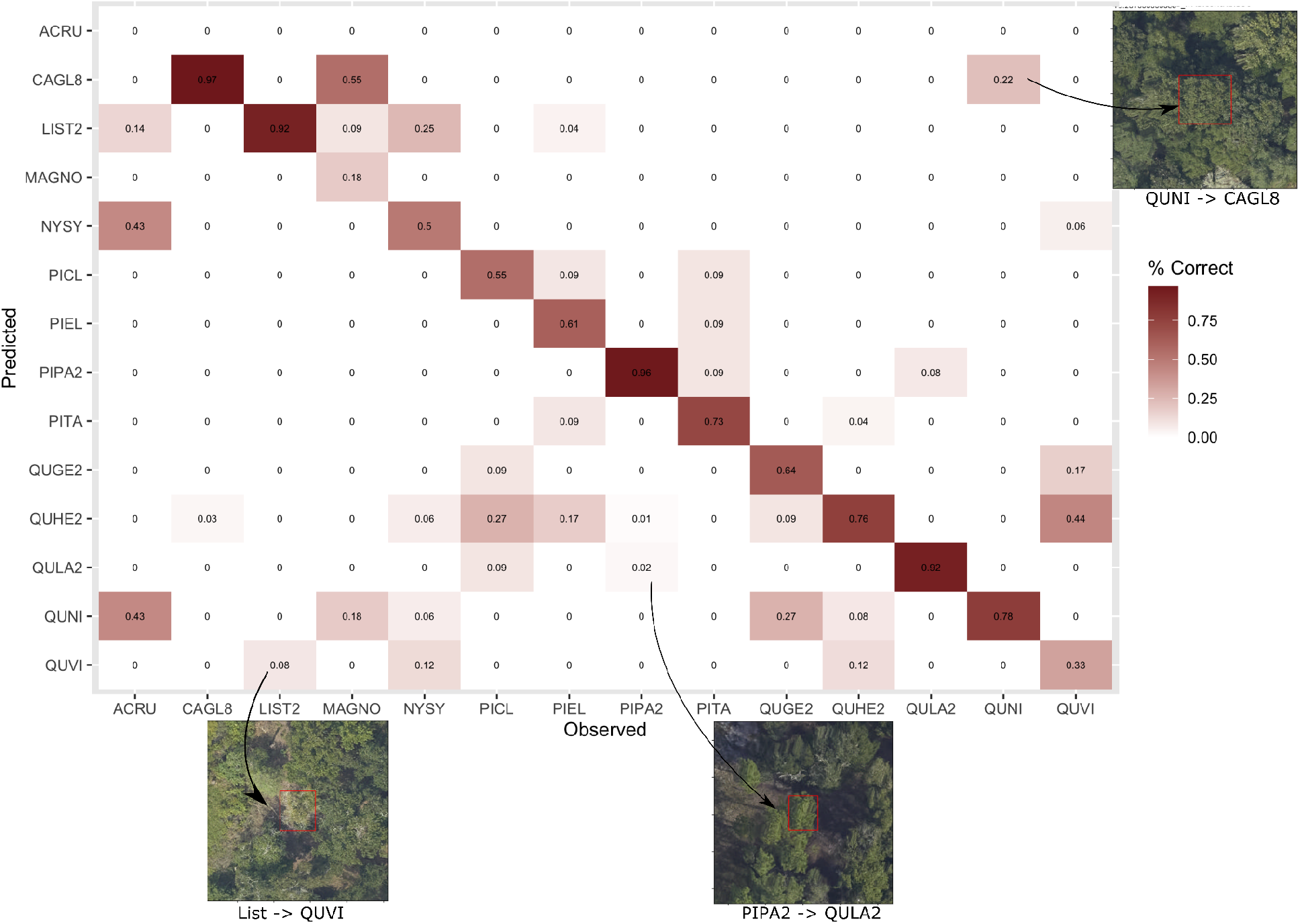
Confusion matrix between the predicted species for the model using multi-temporal mixture of experts and the field labeled taxa. The proportion of data in each cell is shown and colored.

We predicted the locations of all standing trees within the OSBS area using the existing deepforest RGB model, resulting in 670,883 predicted canopy trees within the OSBS boundary. We filtered these tree crowns using the alive-dead model. The evaluation accuracy of the alive-dead model was 95.8% (Table S1). We identified 22,359 standing dead trees in the landscape. After removing these trees, we classified each tree crown using the multi-temporal hierarchical model. Creating spatial maps of predicted tree crowns showed significant micro-habitat specialization among species. *P. palustris* and *Q. laevis* are the dominant species in the upland, sandhill areas of the site. The habitats at lower elevation and closer to the lakes, particularly xeric hammock and transitional hardwood sites, are dominated by *Q. hemisphaerica and Q. geminata*. An increase in rarer broadleaf species such *Liquidambar styraciflua* and *Nyssa sylvatica* in mesic hammock and baygall habitats, particularly in the southwest portion of the site (Figure 4, Figure S1). There was a low confidence area in the southwest which corresponded to the baygall habitat and increased prevalence of rarer broadleaf species, including *N. sylvatica* and *Acer rubrum* (Figure 5, Figure S1). The zero evaluation accuracy of ACRU should be taken with some consideration given that there are only 6 available evaluation samples, the lowest of any species.

**Figure 4.**
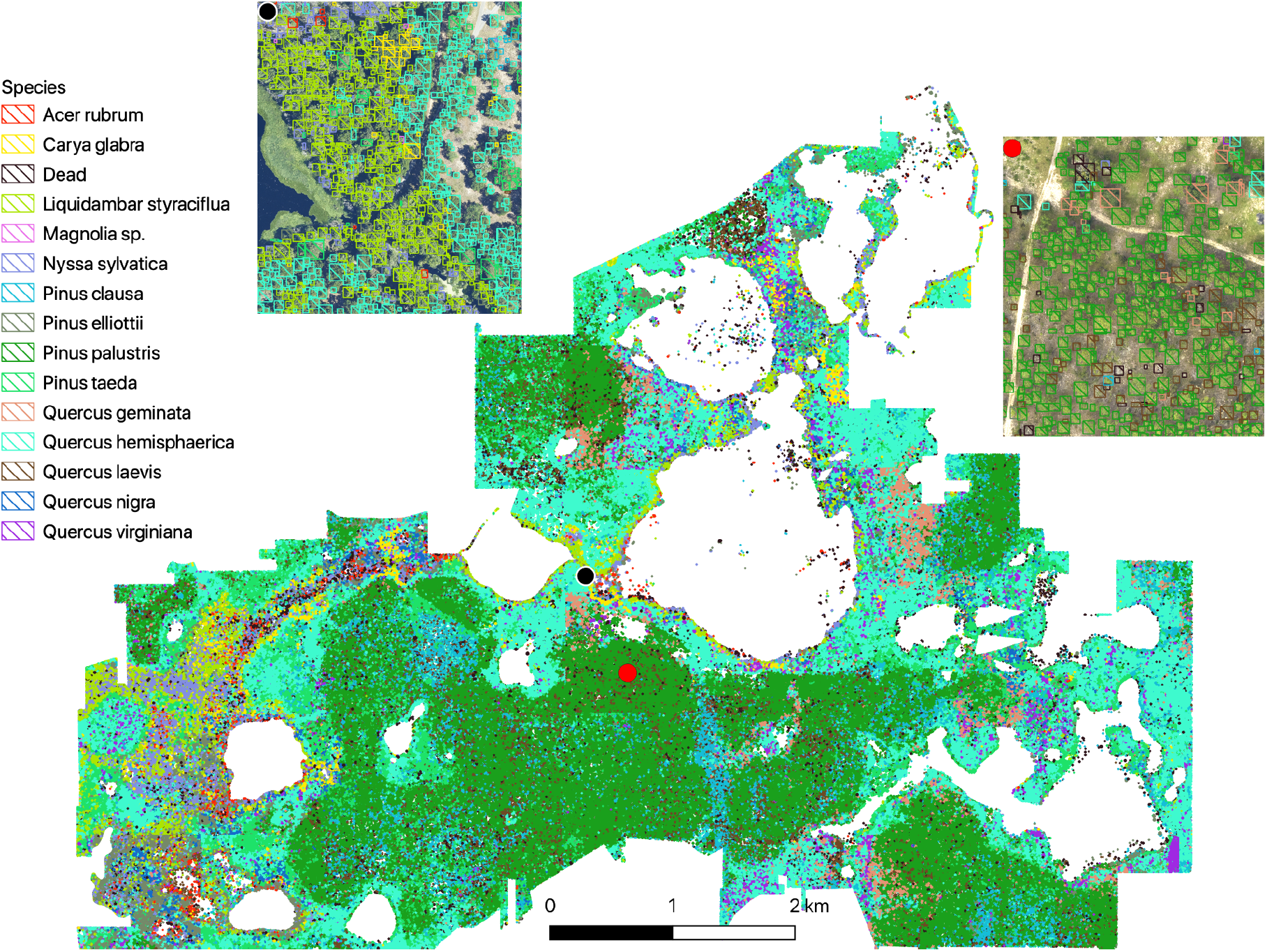
Predicted crown map for 14 species and dead trees at the full site scale (670,883 trees) for the Ordway-Swisher Biological Station. The ensemble species prediction is shown as the result of a five layer hierarchical multi-temporal model. A 100m inset is shown from a biodiverse region of the site, showing where a lake ecosystem meets a hardwood forest (black point) and uphill pine savanna (red point).

**Figure 5.**
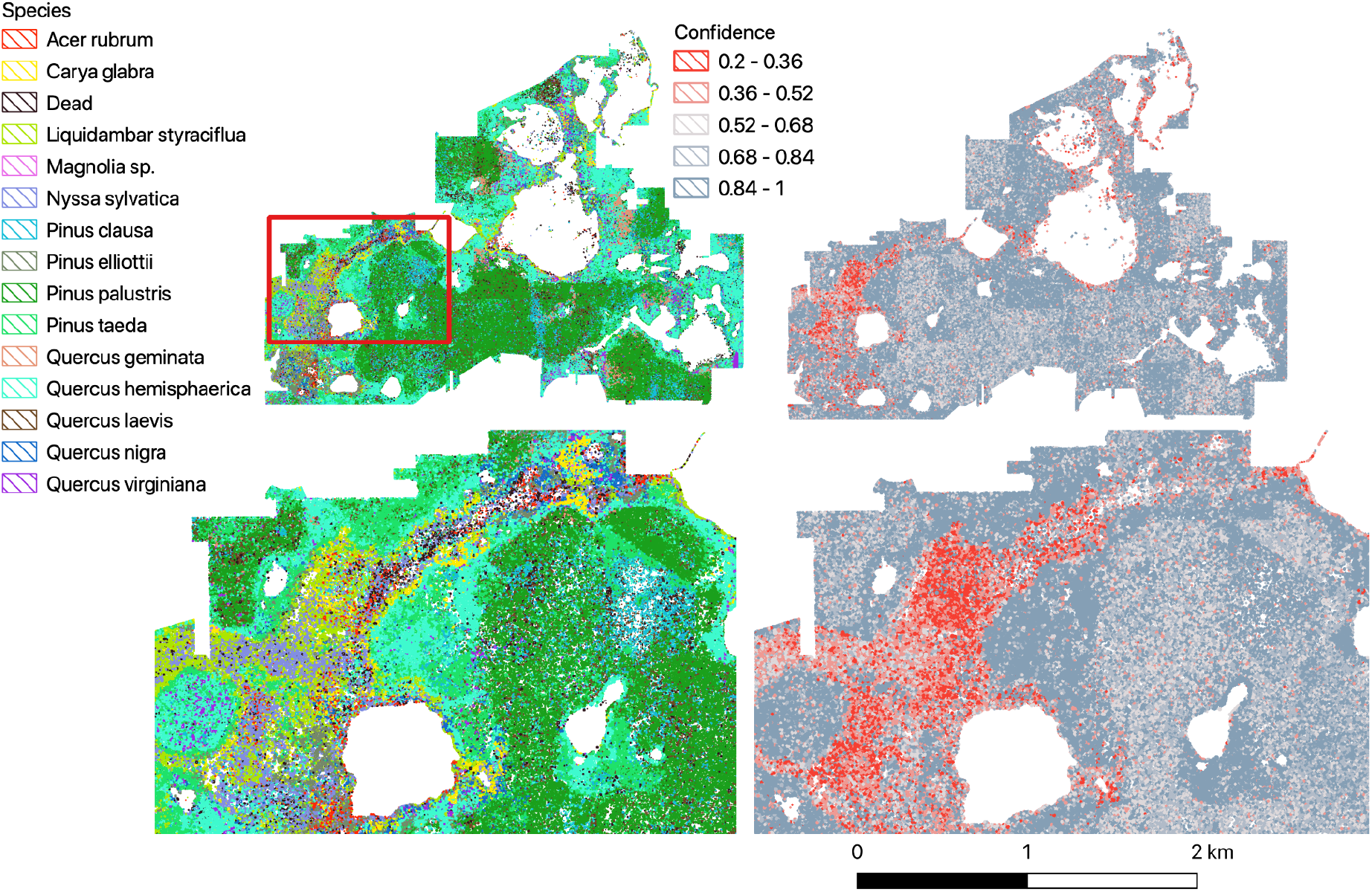
Spatial map of ensemble confidence scores alongside the predicted classes. The confidence score for each predicted species label is shown with low confidence samples shown in red and high confidence in blue.

## Discussion

Using multi-year airborne hyperspectral data, an expanded set of field-labeled tree stems focused on sampling rare species and habitats, and computer vision approaches designed to support classification of rare species, we produced individual level crown maps for 14 tree species and a dead class at the Ordway-Swisher Biological Station (OSBS) NEON site with 75% accuracy. These 14 species represent almost 90% of all stems present at the site and include rare species representing less than 1% of the individual trees on the landscape. Compared to the single-site OSBS model created from NEON woody vegetation structure data alone (Marconi et al. 2022, evaluation accuracy in Table S2), we doubled the species number predicted by incorporating auxiliary, non-NEON data. As a result, 25% of crowns predicted at OSBS in this study were of species not included in previous efforts. This means that earlier predictions had a ceiling of 75% accuracy for full site predictions, regardless of their performance. This highlights the limitations of fixed forest inventory plot data to develop remote sensing models and, as a result, the difficulty of measuring tree biodiversity across large landscapes.

Using targeted data collection alongside NEON field plots, we combined multi-temporal classification models with hierarchical models to improve overall accuracy from 0.61 to 0.75 compared to a baseline CNN model. This approach was particularly effective at improving prediction accuracy of rarer species; average species accuracy improved from 0.48 to 0.63 in the final model. The multi-temporal and hierarchical model complimented each other, with the combined model improving the overall performance more than any component part. Model improvement for rare species did not come at the expense of reduced precision of the dominant *P. palustris* class, which had a recall and precision of 0.96. The improvements of the multi-temporal approach are likely due to the stabilized model learning that comes from multiple lighting conditions that control differences in spectral reflection or image alignment. The drawback of the multi-temporal approach is that it requires the interyear georectification to be consistent. NEON airborne remote sensing surveys are designed to occur at the same time each year (September for OSBS), meaning that images do not capture differences in species phenology (Xi et al. 2021, Veras et al. 2022). Our results suggest that multiple temporal views, even absent phenological information, can benefit generalization when dealing with low training sample sizes (Takahashi Miyoshi et al. 2020, Beery et al. 2022).

The hierarchical model organization contributed to improved rare accuracy of species classification by separately modeling common species and then grouping remaining taxa by general phylogeny and habitat. This approach limits the need for oversampling rare classes or undersampling dominant classes. It may also act as a form of regularization against mislabeled data or samples that are polluted with pixels from neighboring species. By separating classes, we minimized the need for tradeoffs in learning features that separate species classes and allowed each model to learn features more closely refined for smaller target taxa. While this strategy works well when combined with the multi-temporal ensemble, yielding an improvement in micro accuracy of more than 10%, creating, training, and validating hierarchical models is time-consuming and requires additional computational and developer time. Finding an automated way to split classes into nested models will be key in scaling to hundreds of species classes (Liu et al. 2019).

An important methodological change from previous tree species classification workflows is that we generated crown predictions for evaluation using an RGB crown detection model independent from the field labeled trees.This approach reflects the process during prediction in which field stem points are not available. The majority of previous papers use fixed size boxes around field stem points or hand-drawn crown polygons on the imagery to generate both training and evaluation data (Maschler et al. 2018, Scholl et al. 2020, Onishi et al. 2022, Marconi et al. 2022). This crown delineation approach biases results towards higher accuracy as field points are most often collected on the largest and easiest to differentiate individuals. This in turn biases the evaluation score compared to large scale predictions, since during prediction we cannot hand delineate crowns. The downside of using algorithmically generated bounding boxes is that it introduces additional uncertainty due to imperfect crown predictions. The dominant species, *P. palustris*, happens to be relatively easy to delineate due to regular spacing among individuals and simple crown geometry. In contrast, the *Quercus* species have complex crowns and field assessments suggest that the predicted crowns are sometimes oversegmented, leading to multiple predicted crowns for each true tree. This means that the *Quercus* individual counts are biased to be higher compared to the *Pinus* counts. Cascading uncertainty from the crown detection into the species classification model could assist in creating more accurate counts that reflect the bias among species in crown detection accuracy (Maschler et al. 2018).

We applied the resulting model to make landscape maps capturing 670,883 individual trees from 14 species, plus a dead tree class, at OSBS. Using the uncertainty in data measured by cross-validation (in red, Figure 6), we estimate there are 189,878 *P. palustris* trees (lower 5th quantile=147,240, upper 5th quantile=226,880), which equates to 28.3% (21.9%, 33.8%) of the detected trees. *P. palustris* mostly occurs in the higher elevation sandhill pine forests co-occurring with *Q. laevis* and to a lesser extent *Q. geminata*, and *Q. hemisphaerica* in the hardwood forests at lower elevation and fringing lakes alongside other broadleaf species. The remaining species each represented less than 10% of crowns, with 10 species below 5% of crowns. The rarest two species, *A. rubrum* and *Magnolia sp*. were predicted to be less than one tenth of 1% of crowns. There is a higher concentration of rare broadleaf species in the southwest area of the site, corresponding to a low lying “baygall” swamp ecosystem (Figure 5, Figure S1). We have since visited these areas and confirmed the tree biodiversity contains a higher proportion of *N. sylvatica* than the rest of the site. However, these predictions are likely overly biased towards this species and are a proxy for the higher tree diversity in the swamp rather than a single patch composed dominantly of *N. sylvatica*. The low coverage of training data in this area likely led to the model predicting that the area was dominated by one class that was collected nearby. This example underscores the complexity of assessing diversity in broad scale forested ecosystems and the virtue of interactive sampling efforts where areas with low prediction confidence merit additional data collection.

**Figure 6.**
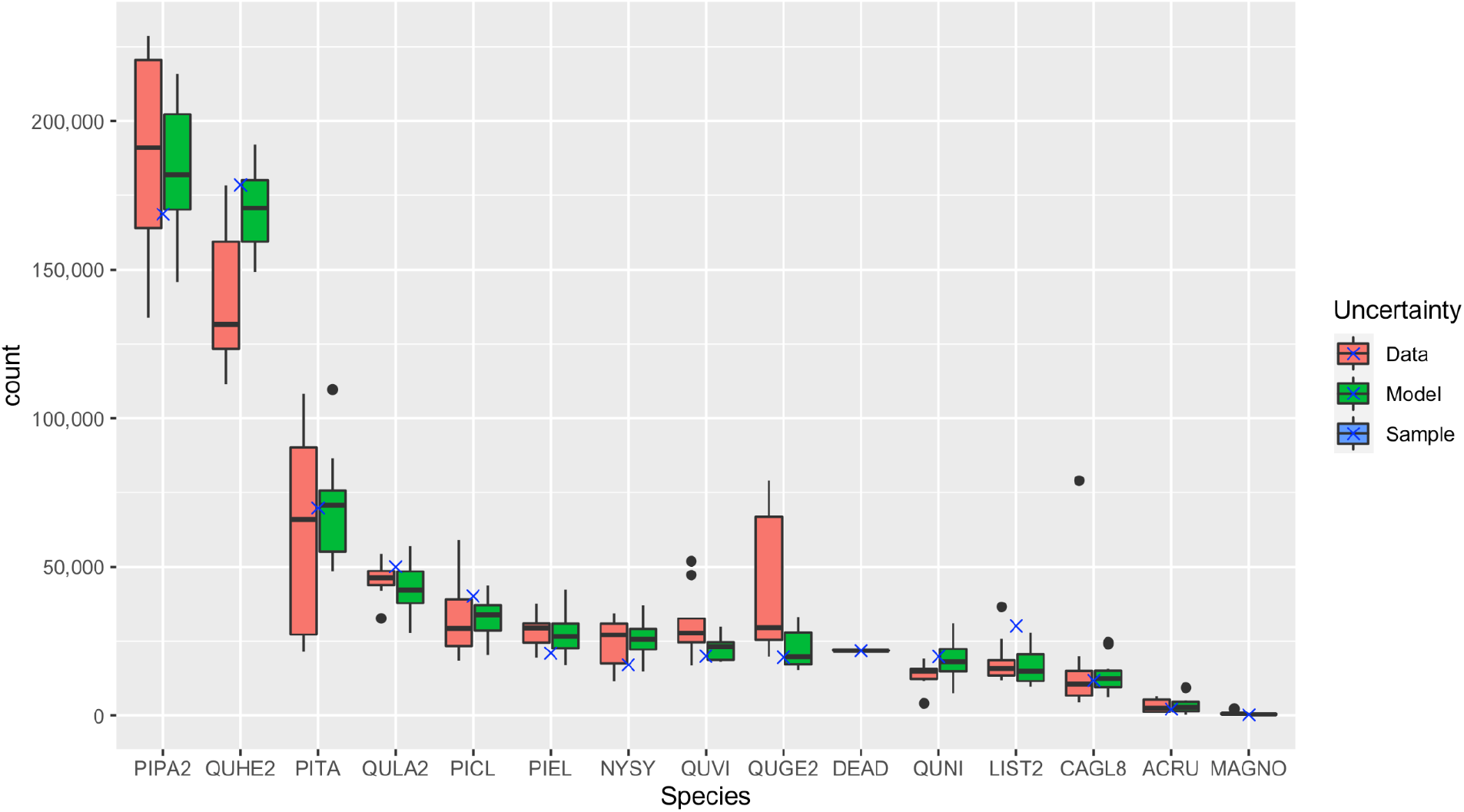
Uncertainty in predicted counts for the entire OSBS NEON site using the multi-temporal hierarchical model. The ‘Data’ uncertainty is assessed using cross-validation of the train/test splits from the original field collected data. For each train/test split, the entire workflow is run to create predictions of species abundances at the site-level. The ‘Model’ uncertainty is assessed using repeat model training for the same train/test split. This uncertainty reflects the stochastic nature of the model optimization process. The error bars on both types of uncertainty are the result of predictions across 10 iterations. The ‘Sample’ uncertainty, represented by an “X”, is a multinomial draw from the evaluation confusion matrix based on the confidence score of each predicted tree crown.

From these spatially explicit predictions, we aggregated the data to estimate the total number of individuals of each species at the site. Given the low number of training and testing samples for rare species, estimating the uncertainty in predicted counts is crucial for conducting downstream ecological analysis using remote-sensing derived data. This has been generally overlooked in previous work due to ongoing debate about how to interpret machine learning outputs and translate them into measures familiar to ecologists. We offered three measures of uncertainty that gave rise to reasonably similar numbers of counts, with little change in the rank abundance among taxa. The exception was the tradeoff between *Q. hemisphaerica* and *Q. geminata*. These two species are closely related and have only modest habitat differences. Overall, the landscape maps do a good job at showing the general pattern of oak ecology, with Q. *hemisphaerica* more common in upland areas, *Q. geminata* more common in the sandy dry areas, and *Q. virginiana* in more mesic hardwood ecosystems. These species are known to hybridize, and the higher uncertainty in *Q. geminata* counts underscores the importance of assessing models using train-test cross validation when predicting across large landscapes.

The three types of uncertainty used in this paper provide a first glimpse into the integration between computer vision and ecological modeling. In particular, the per-sample uncertainty is a coarse measure since neural networks tend to be overconfident, leading to reduced predicted uncertainty (Abdar et al. 2021). The approach of multiplying each sample by the confusion matrix is complicated by the lack of granularity in the evaluation data. If there are only 5 evaluation samples for a particular taxa, then the evaluation score is quantized to come in sizes of 20%. When assessing trees at the scale of millions of examples, this lack of resolution makes the calculation sensitive to idiosyncratic examples that could lead to erroneous allocation of predicted counts. For this reason, forcing the inclusion of species with a low number of evaluation samples can add confusion to the model without reliable information on prediction accuracy. For example, while the *Acer rubrum* evaluation accuracy in Figure 3 is 0%, we believe it is very unlikely that all *A. rubrum* predictions in landscape maps are incorrect. We simply lack the evaluation data (n=6) to assess its true accuracy. Deeper integration between deep learning neural networks and traditional hierarchical Bayesian models will help clarify the relationship between the predicted class conditional on the sample probability and the prior expectation of confusion among species.

The availability of large scale remote sensing has opened the potential for monitoring and analyzing ecological data at unprecedented scales. However, while ecological research often requires information about rare species, many remote-sensing based approaches are forced to ignore those rarer classes due to limitations in data availability and training, evaluating and model design. We have shown that for a single site it is possible to increase the coverage of biodiversity in remote sensing models by leveraging additional sources of data, targeting sampling of rarer classes, using multiple images of each individual, and employing hierarchical model architectures. Deciding on the level of rarity captured by remote-sensing models that is necessary for conducting ecological studies remains an important question for researchers moving forward.

## Data Availability

The github repo has been archived on zenodo https://github.com/weecology/DeepTreeAttention (https://zenodo.org/record/7308745#.Y3KiwOzMKHE) along with hyperspectral crops from the train and test split used to generate the landscape maps https://zenodo.org/record/7301868. The RGB model used to generate predicted tree crowns is available through the deepforest python package: https://deepforest.readthedocs.io/.

## Acknowledgements

We would like to thank NEON staff and in particular Tristan Goulden and Courtney Meier for their assistance and support. We thank Natalie Heaton, Nicollete Lyons, Matthew Raulerson, Alex Seeley, Camille Sicangco, Luis Tirado and Stuart Wilkin and for field data collection efforts. This research was supported by the Gordon and Betty Moore Foundation’s Data-Driven Discovery Initiative (GBMF4563) to EP White, by the USDA National Institute of Food and Agriculture McIntire Stennis project 1024612 to SA Bohlman, and by the National Science Foundation (1926542) to EP White, SA Bohlman, A Zare, DZ Wang, and A Singh.

## Author Contributions

EW, SB, AZ, AS, SM, SG conceived the study; SM, PT, DJ, and LM collected data, BW and SM performed the analysis. All authors contributed to manuscript writing and figure creation.

## Supplemental Figures

**Figure S1.**
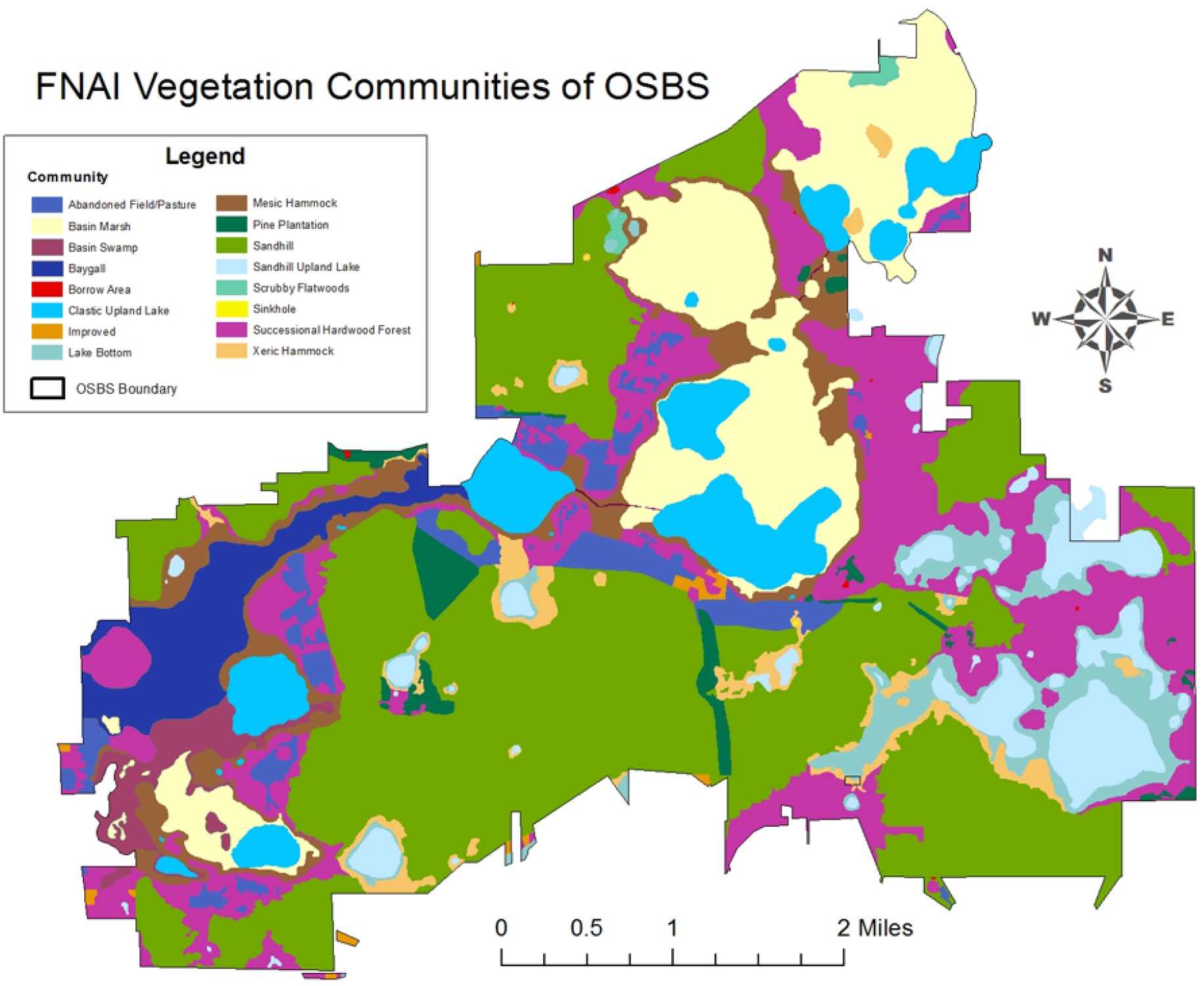
Florida Natural Areas Inventory map for the OSBS site. Obtained from fnai.org.

**Table S1.** Confusion matrix for the Alive/Dead model to filter out predicted trees before species classification using the hyperspectral model.

|  |  | Predicted |  |
| --- | --- | --- | --- |
|  |  | Alive | Dead |
| Observed | Alive | 527 | 9 |
|  | Dead | 10 | 89 |

**Table S2.**
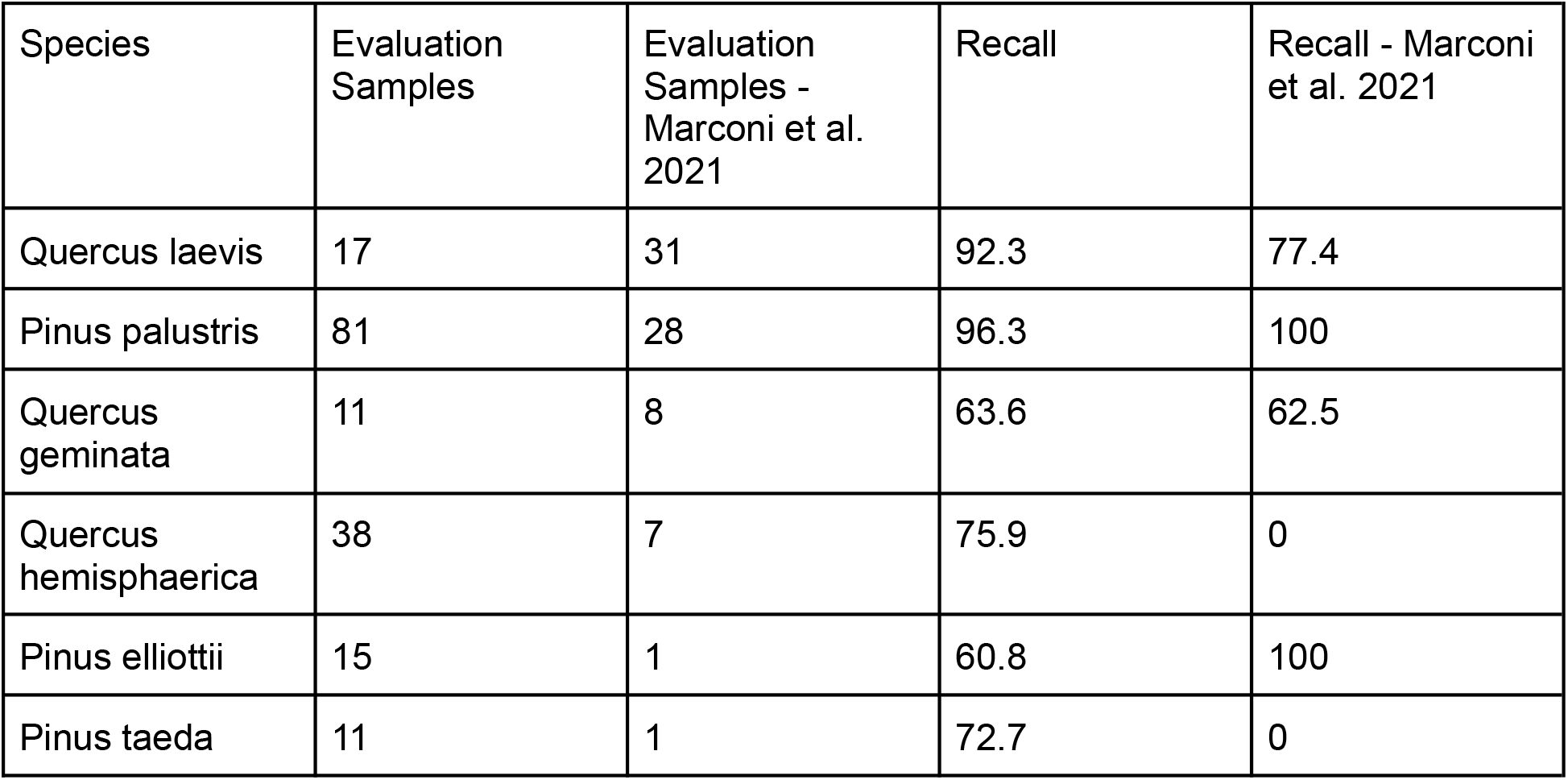
Species results for the six species that occur in Marconi et al. 2021. Sorted by number of evaluation samples in Marconi et al. 2021.

| Species | Evaluation Samples | Evaluation Samples - Marconi et al. 2021 | Recall | Recall - Marconi et al. 2021 |
| --- | --- | --- | --- | --- |
| Quercus laevis | 17 | 31 | 92.3 | 77.4 |
| Pinus palustris | 81 | 28 | 96.3 | 100 |
| Quercus geminata | 11 | 8 | 63.6 | 62.5 |
| Quercus hemisphaerica | 38 | 7 | 75.9 | 0 |
| Pinus elliotii | 15 | 1 | 60.8 | 100 |
| Pinus taeda | 11 | 1 | 72.7 | 0 |

